# Spatiotemporal specificity of correlated DNA methylation and gene expression pairs across different human tissues and stages of brain development

**DOI:** 10.1101/2021.05.12.443776

**Authors:** Kangli Wang, Rujia Dai, Yan Xia, Jianghua Tian, Chuan Jiao, Tatiana Mikhailova, Chunling Zhang, Chao Chen, Chunyu Liu

**Affiliations:** Center for Medical Genetics & Hunan Key Laboratory of Medical Genetics, School of Life Sciences, and Department of Psychiatry, The Second Xiangya Hospital, Central South University, Changsha, Hunan, China; Department of Psychiatry, State University of New York Upstate Medical University, Syracuse, NY, USA; Department of Neuroscience and Physiology, State University of New York Upstate Medical University, Syracuse, NY, USA; National Clinical Research Center on Mental Disorders, The Second Xiangya Hospital, Central South University, Changsha, Hunan, China; National Clinical Research Center for Geriatric Disorders, Xiangya Hospital, Central South University, Changsha, Hunan, China

**Keywords:** DNA methylation, gene expression, tissue, development, gene and CpG pairs (GCPs)

## Abstract

DNA methylation (DNAm) that occurs on promoter regions is primarily considered to repress gene expression. Previous studies indicated that DNAm could also show positive correlations with gene expression. Both DNAm and gene expression profiles are known to be tissue- and development-specific. This study aims to investigate how DNAm and gene expression are coordinated across different human tissues and developmental stages, as well as the biological significance of such correlations. By analyzing 2,239 samples with both DNAm and gene expression data in the same human subjects obtained from six published datasets, we evaluated the correlations between gene and CpG pairs (GCPs) at cis-regions and compared significantly correlated GCPs (cGCPs) across different tissues and brains at different age groups. A total of 37,363 cGCPs were identified in the six datasets; approximately 38% of the cGCPs were positively correlated. The majority (>90%) of cGCPs were tissue- or development-specific. We also observed that the correlation direction can be opposite in different tissues and ages. Further analysis highlighted the importance of cGCPs for their cellular functions and potential roles in complex traits and human diseases. For instance, early developmental brain possessed a highly unique set of cGCPs that were associated with neurogenesis and psychiatric disorders. By assessing the epigenetic factors involved in cGCPs, we discovered novel regulatory mechanisms of positive cGCPs distinct from negative cGCPs, which were related to multiple factors, such as H3K27me3, CTCF, and JARD2. The catalog of cGCPs compiled can be used to guide functional interpretation of genetic and epigenetic studies.

## Introduction

DNA methylation (DNAm) at CpG sites is a major epigenetic marker that regulates gene expression and thus participates in many essential biological processes and human diseases(Reik 2007; Barlow 2011; Baylin and Jones 2011; Jones 2012). DNAm is commonly understood to suppress gene expression by blocking the binding sites of transcription factors at gene promoter regions(Jaenisch and Bird 2003; Stadler et al. 2011). However, studies revealed that DNAm in the gene body could be positively correlated with gene expression(Hellman and Chess 2007; Lister et al. 2009). Genome-wide evaluation of DNAm-expression associations also supported the presence of positive correlations(Olsson et al. 2014; Wagner et al. 2014; Tasaki et al. 2018; Taylor et al. 2019). There is multiple evidence suggesting a spatiotemporal relationship between DNAm and gene expression. DNAm is reported to be development- or tissue-specific, and correlated with the expression of corresponding genes(Liang et al. 2011; Bell et al. 2012; Smith and Meissner 2013; Varley et al. 2013; Reynolds et al. 2014; Blake et al. 2020). Comparison of methylation patterns in fibroblasts, T-cells, and lymphoblastoid cells suggested that differentially methylated regions during development could contribute to delineating gene expression in different cell types(Gutierrez-Arcelus et al. 2013; Gutierrez-Arcelus et al. 2015). Bonder et al.(Bonder et al. 2014) assessed this correlation by analyzing 158 liver samples from two studies and compared the liver to the muscle and adipose tissues. Their results indicated more negative DNAm-expression correlations than positive correlations and the correlations showed strong tissue-specificity. However, the developmental specificity of the DNAm-expression relationships has not been studied to date. Moreover, these studies did not examine the implications of DNAm in cellular functions nor the connection between DNAm-expression correlations and human diseases.

The DNAm-expression relationship is affected by numerous biological factors. The relative location of CpG within the gene body may influence regulation of expression and thus DNAm-gene expression relationships. It has been reported that CpGs on promoter regions are negatively correlated with gene expression while those on gene bodies function quite the opposite(Jones 2012). DNAm on CpG islands (CGI) has been found to be associated with transcriptional repression and gene silencing(Jones 2012; Schubeler 2015). In addition, some transcription factors (TFs) such as RFX5 preferred DNAm and could interact with methylated-DNA, while some others such as NRF1 have been reported to be more sensitive to DNAm and only bind unmethylated sequences, thereby affecting the gene expression(Schubeler 2015; Zhu et al. 2016; Yin et al. 2017; Wang et al. 2018). A comprehensive epigenomic study revealed that chromatin state could also influence DNAm and gene expression(Roadmap Epigenomics et al. 2015). Accessible chromatin, marked by promoter DNAm depletion, histone acetylation, and H3K4 methylation, is believed to facilitate transcription(Collings and Anderson 2017; Lovkvist et al. 2017; Spektor et al. 2019). In contrast, histone modification such as H3K9me3 is linked to the establishment of DNAm and is involved in heterochromatin and long-term gene repression(Cedar and Bergman 2009; Du et al. 2015). Furthermore, a study of blood found that CpGs negatively correlated with gene expression are enriched in transcriptionally active regions such as enhancers while CpGs positively correlated with gene expression occur in repressed regions such as Polycomb-repressed regions(Bonder et al. 2017). Chromatin structure alterations can also change the relationship between DNAm and gene expression(Liu et al. 2016). Taken together, numerous factors might influence DNAm-gene expression relationships. Yet the exact epigenetic mechanisms of the positive or negative correlation of DNAm-expression remain elusive.

In this study, we intend to confirm the spatiotemporal specificity of the correlated relationships between DNAm and gene expression in multiple tissues, particularly brain tissue that showed highly dynamic changes of gene expression and DNAm during development(Kang et al. 2011; Numata et al. 2012), and to explore its biological significance as related to disease association, and to identify epigenomic features related to positively and negatively correlated DNAm-expression relationships. Spatial specificity refers to DNAm-expression correlations that can only be detected in specific tissues but not the others, while temporal specificity refers to correlated relationships that change with development and aging processes. We constructed gene and CpG pairs (GCPs) using gene expression and DNAm data from the same human subject available in six public datasets (**Fig.1A**), and further compared significantly correlated GCPs (cGCPs) firstly across a group of different tissues (brain, liver, monocytes, and T-cells), and secondly across exclusively brain tissue at three different age groups (“developmental brain” with age ranging from fetus to age 25 years old (y.o.); “aging brain” with age of 25 y.o. to 78 y.o.; and “aged brain” from all senior donors with age at 88±7 y.o. and limited age variation). Gene functions and disease associations of these cGCPs were subsequently analyzed. Finally, we investigated the enrichment of epigenomic features on both positively and negatively correlated GCPs to identify the potential driver of opposite correlations. Our genome-wide analysis revealed unique DNAm-expression profiles in different tissues and stages of brain development consistent with expected tissue-specific functions and developmental processes.

**Figure 1.**
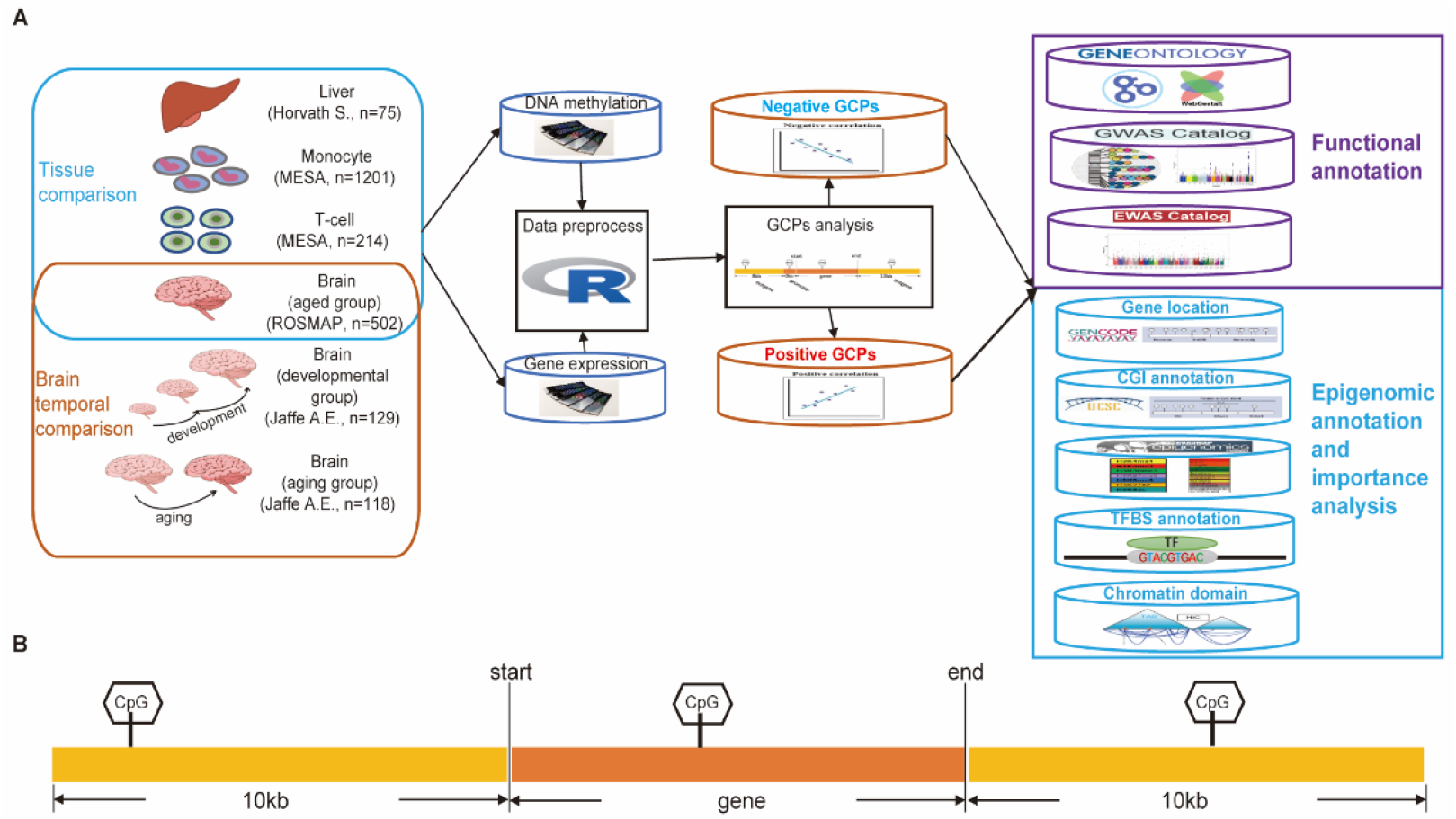
The Overview of study design. (A) Design of the study. (B) Gene and CpG Pairs (GCPs) are defined as CpG sites located within the 10kb flanking region of the corresponding gene.

## Results

### 1. More than 30% of the cGCPs were positively correlated in different tissues and developmental brain

We collected six datasets that have both methylation and matched expression data. After quality evaluation and filtering, we retained the data of 502 brains (also referred to as brain aged group), 75 livers, 214 T-cell, 1201 monocyte samples from adult human subjects, as well as 129 human brains from donors that were fetus to 25 years old, referred to as the brain developmental group, and 118 human brains from donors that were 25 years old to 78 years old, referred to as the brain aging group (**Supplemental Table S1, Fig.1A**). DNA methylation measurements were performed using Illumina HumanMethylation450 BeadChips to interrogate more than 485,000 CpG sites in all datasets. CpGs with variations under 50% of the probes were filtered out based on the median absolute deviation (MAD) of DNAm levels to focus GCP analysis on highly variable loci. The numbers of CpGs and genes used in the analysis are shown in **Table 1** (criteria described in Methods).

**Table 1.** Summary of significantly correlated GCPs (BH adjusted p-value < 0.05) in each tissue. GCPs, gene and CpG pairs; cGCPs, correlated GCPs; ncGCPs, negatively correlated GCPs; pcGCPs, positively correlated GCPs.

| Tissues | # of paired samples | # of total methylation probes after preprocess | # of total expression probes after preprocess | # of total GCPs | # of cGCPs | # of ncGCPs | Mean coeff. of ncGCPs | # of pcGCPs | Mean coeff. of pcGCPs | pcGCPs rate |
| --- | --- | --- | --- | --- | --- | --- | --- | --- | --- | --- |
| Brain DLPFC<br>(also as brain aged group, ROSMAP) | 502 | 214,611 | 17,026 | 230,087 | 8,828 | 5,530 | -0.22 | 3,298 | 0.19 | 0.37 |
| T-cells<br>(Reynolds L.M.) | 214 | 221,838 | 11,498 | 48,210 | 4,359 | 2,999 | -0.31 | 1,360 | 0.27 | 0.31 |
| Liver (Horvath S.) | 75 | 223,578 | 17,685 | 210,025 | 2,281 | 1,421 | -0.47 | 860 | 0.47 | 0.38 |
| Monocytes (Reynolds L.M.) | 1,201 | 217,681 | 10,953 | 46,831 | 5,134 | 3,157 | -0.17 | 1,977 | 0.15 | 0.39 |
| Brain DLPFC developmental group (Jaffe A.E.) | 129 | 217,677 | 24,198 | 133,901 | 16,500 | 9,697 | -0.32 | 6,803 | 0.32 | 0.41 |
| Brain DLPFC aging group (Jaffe A.E.) | 118 | 217,677 | 24,198 | 133,218 | 261 | 190 | -0.46 | 71 | 0.43 | 0.27 |

GCPs were defined as CpG sites located within the 10kb flanking region of the corresponding gene (**Fig.1B**). As a result, the number of GCPs tested in each tissue varied from 230,087 in the aged brains to 46,831 in monocytes (**Table 1**). Using Spearman’s rank correlation test and the Benjamini-Hochberg (BH) multiple testing correction, we obtained thousands of cGCPs (BH adjusted p-value < 0.05) in different tissues and stages of the developmental brain (**Table 1, Supplemental Table S2**). The largest number of cGCPs (16,500) was obtained from the brain developmental group and the smallest (261) from the brain aging group. Both positively correlated GCPs (pcGCPs) and negatively correlated GCPs (ncGCPs) were detected in each dataset (**Table 1**). The pcGCPs accounted for 27%~41% of GCPs across different tissues and stages of the brain (**Table 1**). Cross- and downsampling-validation of cGCPs supported the robustness of the results described herein (**Supplemental Table S3, Supplemental Fig. S1A, S1B**).

### 2. Majority of the cGCPs were tissue-specific

Comparison across different tissues revealed more than 90% of ncGCPs (10,015/11,125) or pcGCPs (6,243/6,696) to be specific to the liver, T-cells, monocytes, or brain (**Fig. 2A, 2B, Supplemental Fig. S2**). Genes and CpGs in cGCPs were named cGenes and cCpGs, respectively. Then Gene Ontology (GO) functional analysis was performed for the cGenes that harbored tissue-specific cGCPs. Different cGene sets showed significant enrichment for tissue-specific functions in the corresponding tissue (**Fig. 2C**, **Supplemental Table S4-7**). For example, the cGenes in brain were enriched for cell adhesion (BH adjusted p-value = 2.85E-12) in GO biological process (BP) and neuron-specific part (BH adjusted p-value = 2.82E-11) in GO cellular component (CC), innate immune response in monocyte cGenes (BH adjusted p-value = 7.03E-10) in BP, leukocyte activation in T-cell cGenes (BH adjusted p-value = 2.53E-12) in BP, and oxidoreductase activity in liver cGenes (BH adjusted p-value = 3.14E-3) in GO molecular function (MF). The functional annotation of cGenes in cGCPs specific to one tissue was similar to that in total cGCPs in the corresponding tissues (**Supplemental Fig. S3, Supplemental Table S8-11**). This is mainly the result of the tissue-specific character of the cGCPs.

**Figure 2.**
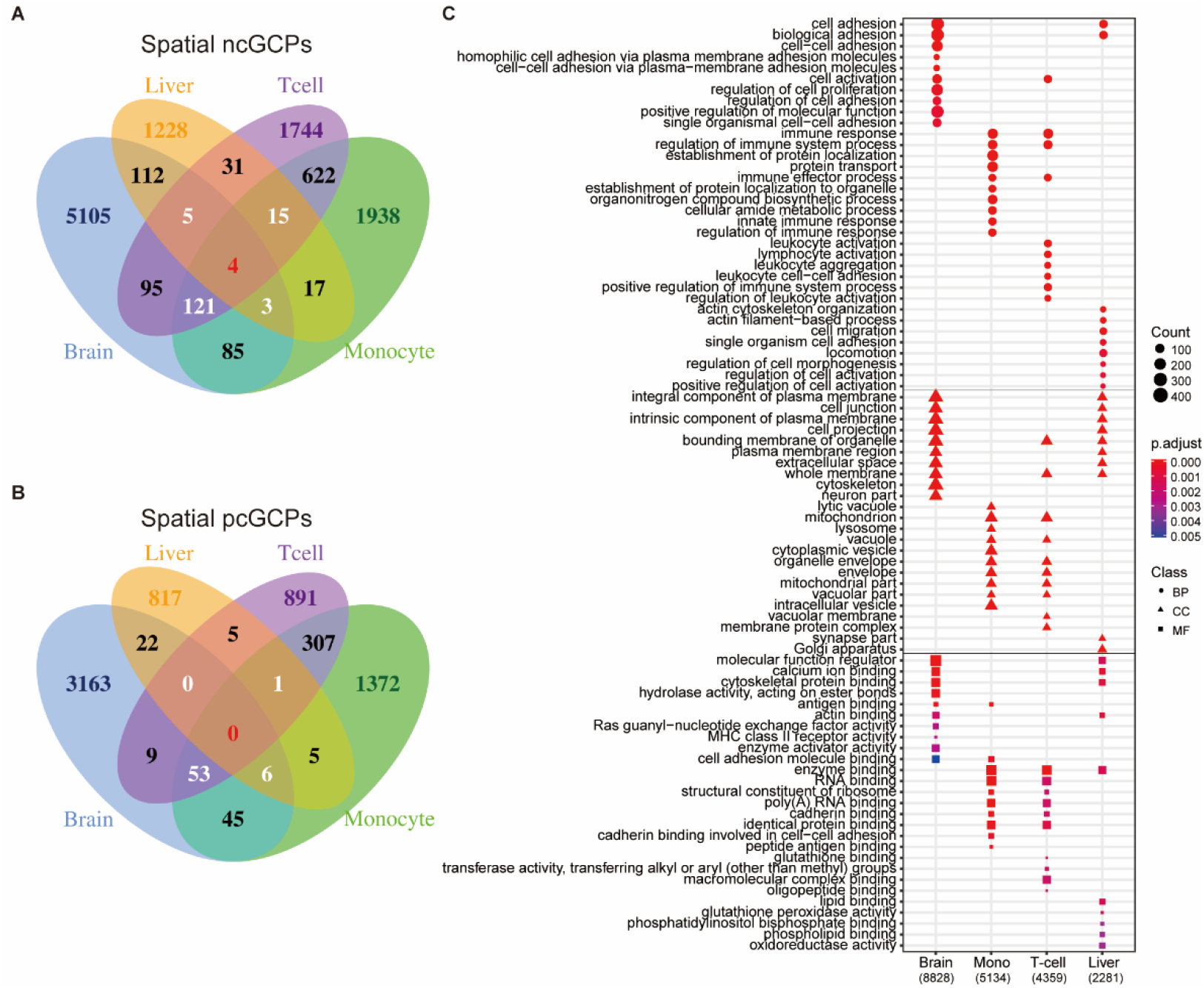
cGCPs are tissue-specific and the cGenes are enriched in tissue-specific functions. (A) The Venn plot to compare negatively correlated GCPs (ncGCPs) across four tissues. Brain, aged brain. (B) The Venn plot to compare positively correlate GCPs (pcGCPs) across four tissues. (C) The top-ranked GO enrichment items for cGenes across four tissues. The number in parenthesis under tissue name refers to cGCPs count. Mono, monocytes.

Even though thousands of cGCPs were identified in different tissues, only 208 of them were shared by three or more tissue types; of which 148 were negatively correlated and 60 were positively correlated (**Fig. 2A, 2B**). Notably, four cGCPs (*CD86*-cg04387658, *GSTM1*-cg10950028, *GSTT1*-cg04234412, *GSTT1*-cg17005068) were shared in all tissues and are consistently negatively correlated (**Fig. 2A, Fig. 3**). The cGenes in the 208 cGCPs shared in at least three tissues were enriched in glutathione derivative metabolic process (BH adjusted p-value = 1.8e-04) in BP, MHC class II protein complex (BH adjusted p-value = 1.13e-06) in CC, and glutathione binding (BH adjusted p-value = 1.36e-04) in MF, which were the top pathways in the three GO categories (**Supplemental Fig. S4, Supplemental Table S12**). *GSTT1* and *GSTM1* in cGCPs shared in four tissues were both related to glutathione derivative metabolic process and glutathione transferase activity.

**Figure 3.**
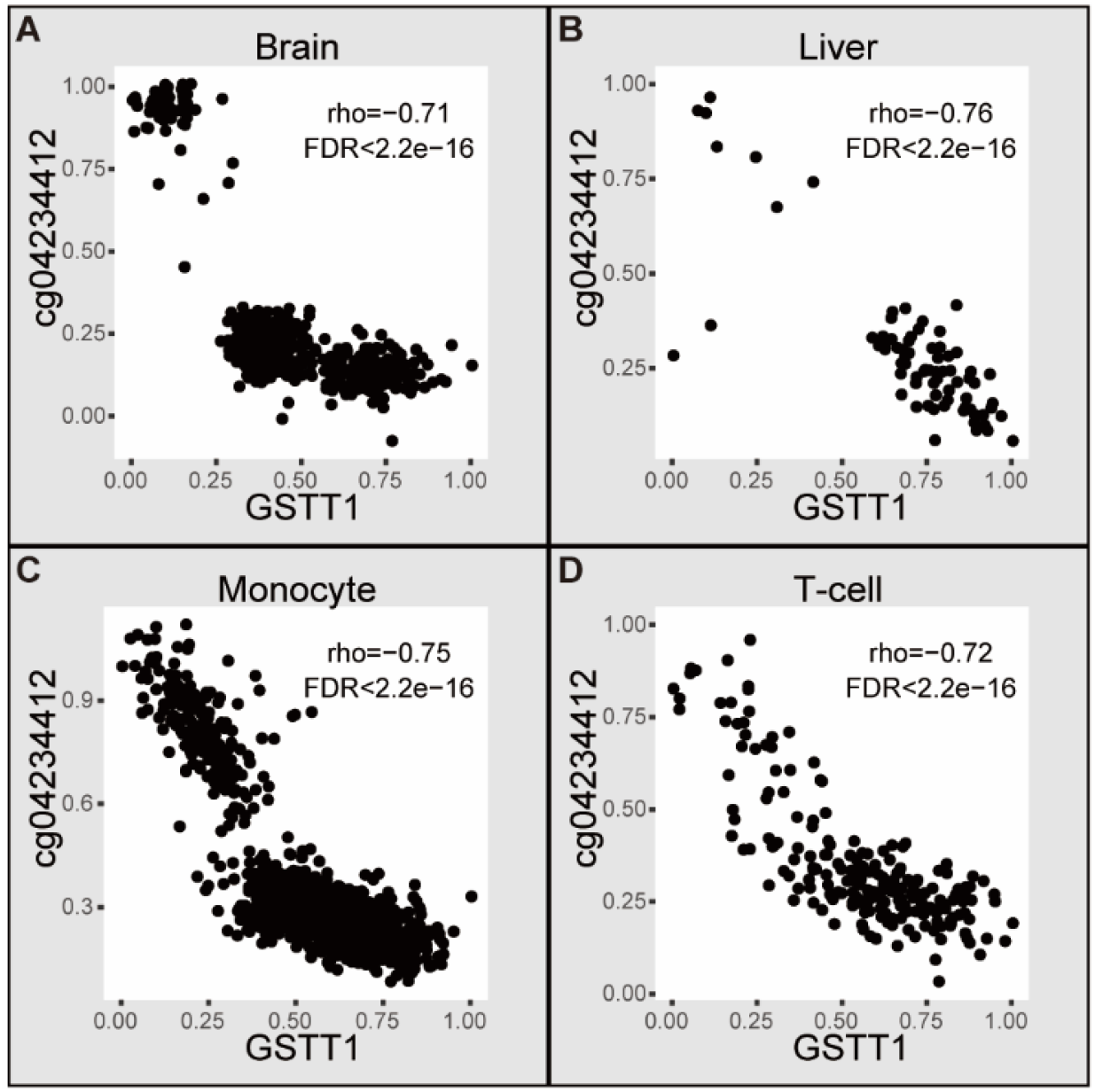
An example of cGCPs shared across four tissues. Scatter plots of *GSTT1* expression level (X-axis) and cg04234412 methylation level (Y-axis) in the aged brain (A), liver (B), monocyte (C), and T-cell (D). Brain, aged brain.

### 3. cGCPs in brain exhibit age specificity

To capture temporal regulation, we examined the age-dependent GCPs throughout the development and aging process of the brain. The brain developmental data were collected from individuals with ages ranging from post-conception (14 weeks) to 25 years old when brain development is complete(Pujol et al. 1993), and the brain aging data were from 25 years old to 78 years old (**Supplemental Table S1**). In brain developmental and aging group 133,901 and 133,218 GCPs were tested and 16,500 and 261 GCPs were significantly correlated, respectively (**Table 1**). Among these cGCPs, 6,803 were pcGCPs in brain developmental group and 71 were pcGCPs in brain aging group.

Furthermore, we compared the cGCPs across the developmental brain, aging brain, and the aged brain groups. The majority (96%) of the cGCPs found in the developmental brain were specific to developmental age group (**Fig. 4A, B**). The cGenes in the developmental brain were uniquely enriched for neurogenesis (BH adjusted p-value < 2.2E-16) and neuron differentiation (BH adjusted p-value < 2.2E-16) pathways in BP that were not present in the other brain groups (**Fig. 4C, Supplemental Table S13**). For the brain aging group, 23% of the cGCPs were specific. No GO terms were enriched for the cGenes of aging brain data. The functional annotation of specific cGenes from the developmental or aged brain was found to be highly analogous to those of the total cGenes in the corresponding brain group (**Supplemental Fig. S5, Supplemental Table S14, S15**), which is most likely due to development-specific cGCPs in the brain tissue.

**Figure 4.**
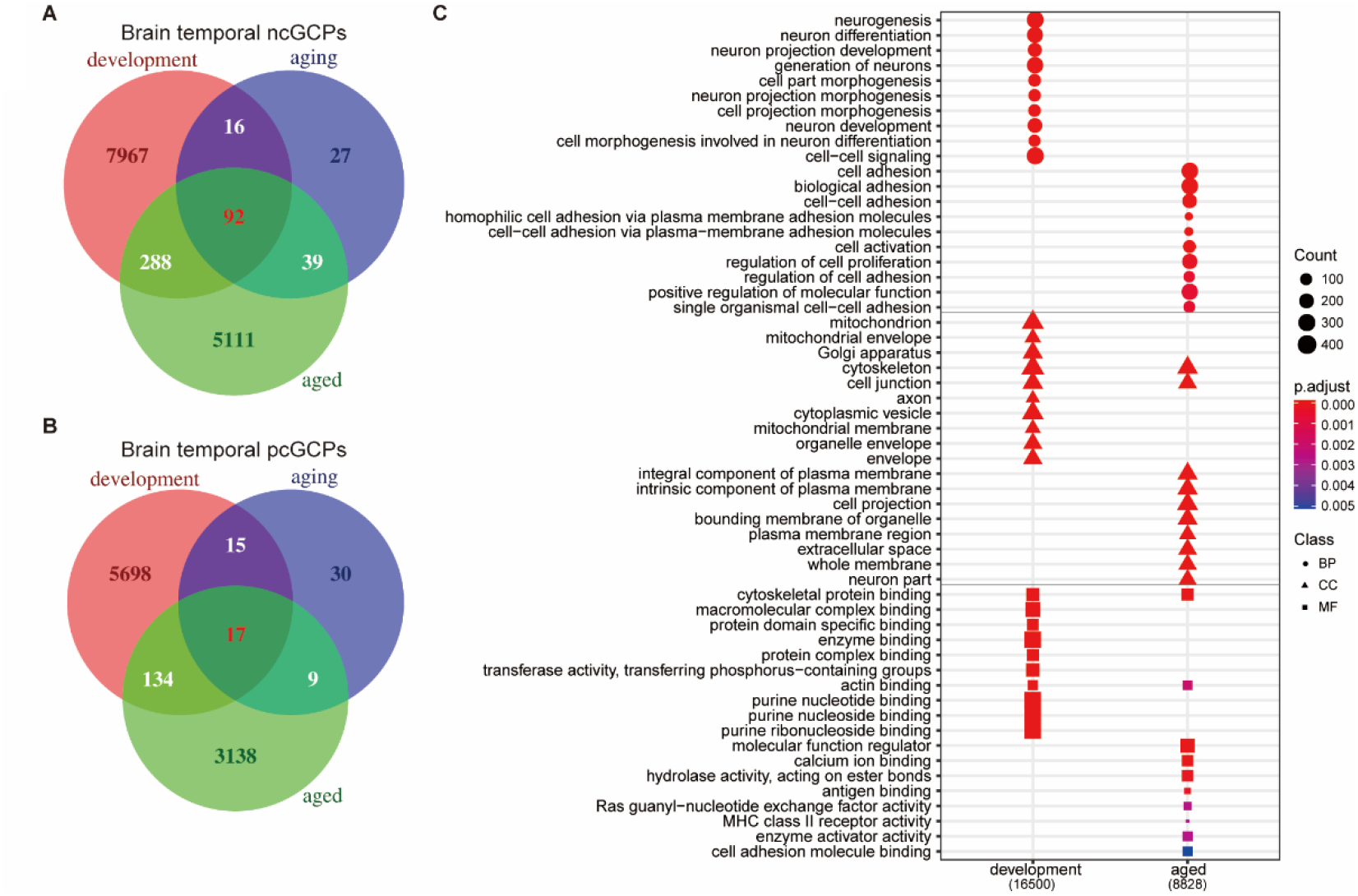
cGCPs in the brain are development-specific and cGenes in the developmental brain are enriched for neurogenesis and neuron differentiation. (A) The Venn plot to compare ncGCPs across the developmental, aging, and aged brain. devBrain, brain developmental group; agingBrain, brain aging group; agedBrain, brain aged group. (B) The Venn plot to compare pcGCPs across the developmental, aging, and aged brain. (C) Comparison of the top-ranked GO enrichment items for cGCPs between the developmental brain and aged brain. The number in parenthesis under stage name refers to cGCPs count.

Additionally, 92 ncGCPs and 17 pcGCPs were shared across three brain age groups (**Fig. 4A, B**). For example, *PCDHB4* gene expression is negatively correlated with cg08626876 methylation level in developmental, aging, and aged brain data (**Supplemental Fig. S6**). The cGenes in these 109 cGCPs were enriched for L-lactate dehydrogenase activity (BH adjusted p-value = 3.16E-02) in MF (**Supplemental Table S16**), which may imply the importance of brain energy use medicated by DNAm.

### 4. The Genes and CpGs of the cGCPs are enriched in traits-relevant tissues

To study the potential role of cGCPs in human diseases, we performed enrichment analysis using genome-wide association study (GWAS) for cGenes and epigenome-wide association study (EWAS) for cCpGs. The enrichments were performed for all phenotypes reported in GWAS and EWAS catalog to avoid potential selection bias.

GWAS enrichment analysis revealed that cGenes of specific tissue types were enriched in GWAS gene sets of the most relevant phenotype. For example, monocyte cGenes were significantly enriched for monocyte count-related gene sets (BH adjusted p-value = 4.08E-05), and liver cGenes were significantly enriched for liver enzyme levels-related gene sets (BH adjusted p-value = 4.93E-02, **Fig. 5A, Supplemental Table S17**). Intriguingly, cGenes in monocytes and T-cells were also enriched for the autism spectrum disorder or schizophrenia-associated genes. Compared to the aging and aged brain, cGenes in the developmental brain were more specifically enriched for genes associated with schizophrenia (BH adjusted p-value = 1.35E-07), bipolar disorder (BH adjusted p-value = 8.61E-04), and educational attainment (BH adjusted p-value = 1.24E-02). Examples of schizophrenia-related cGCPs in three brain groups are shown in **Fig. 5 B-G**. In contrast to *TAP2-cg14812313* pairs correlated in all brain age groups (**Fig. 5 B-D**), *MDK* gene expression is only correlated with cg21009265 methylation level in brain developmental group (**Fig. 5F**).

**Figure 5.**
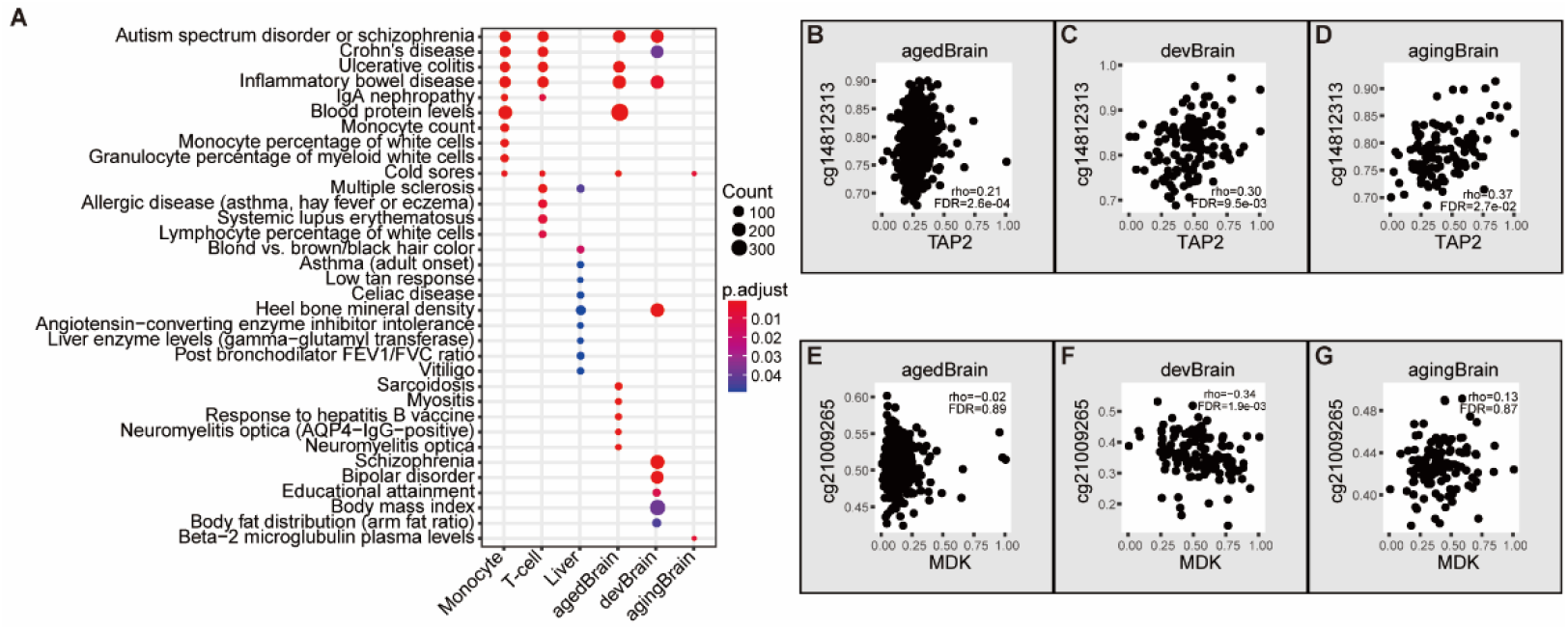
cGenes of specific tissue types are enriched in GWAS of the most relevant phenotype. (A) Top-ranked overrepresented GWAS catalog gene sets in cGCPs across different tissues and brain stages. agedBrain, brain aged group; devBrain, brain developmental group; agingBrain, brain aging group. (B-G) Examples of schizophrenia-related cGCPs in the aged brain developmental brain, and/or aging brain.

Our EWAS enrichment analysis indicated that liver cCpGs were enriched for liver development-related CpGs (BH adjusted p-value = 3.46E-02), and T-cell cCpGs were enriched for the CpGs related to autoantibody production in systemic lupus erythematosus (BH adjusted p-value = 4.65E-02) (**Supplemental Table S18**). In contrast, cCpGs in other tissues were not enriched for phenotype associated CpGs after multiple testing correction.

### 5. Gene expression is correlated both positively and negatively with nearby CpGs, and the correlation direction can be changed

We explored whether expression of specific genes could be correlated with CpGs in both positive and negative manners. In the same data set, an averaged 15.7%±7.4% of genes have been identified in both positive and negative correlations with different CpG sites (**Supplemental Table S19**). For example, in the aged brain, *QSOX2* is negatively correlated with cg14228683 but positively correlated with cg14381623. Furthermore, when considering all the spatiotemporal cGCPs, a gene may be negatively or positively correlated with a CpG site in one data set while showing an opposite correlation with the same or different CpG site in another data set. For instance, *HLA-DQB1* is negatively correlated with cg03202060 in the liver but they show the opposite correlation in T-cells. Among all correlated GCPs across all four tissue types and the developmental and aging brain, up to 36.6% of genes were linked to CpG sites with both positive and negative correlations.

### 6. The ncCpGs and pcCpGs are associated with distinct epigenomic features

To investigate whether mechanisms (termed here as ‘epigenomic features’) associated with gene regulation may influence the DNAm-expression correlation and their direction, we collected various functional features (CGIs, gene location, chromatin states, histone modification, DHS, TF binding regions, and chromatin structures) for CpG annotation. To compare the difference between positive and negative relationships, we performed the enrichment analysis of CpG in ncGCPs and pcGCPs in all data sets. cCpGs in ncGCPs and pcGCPs were named ncCpGs and pcCpGs, respectively.

To assess whether cCpGs location relative to CGI is related to the DNAm-gene expression correlation, we tested the enrichment of cCpGs in CpG island, shore, shelf, and open sea regions. Both pcCpGs and ncCpGs were significantly more likely to be located on CpG island and shore, though they occasionally also lie in the open sea regions (**Fig. 6A**). For example, the ncCpGs and pcCpGs in developmental and aged brain were enriched in CpG island and shore (p-value < 0.05), whereas ncCpGs and pcCpGs in T-cells and ncCpGs in the liver were enriched in open sea regions (p-value < 0.05). However, no instances were found on the CpG shelf.

**Figure 6.**
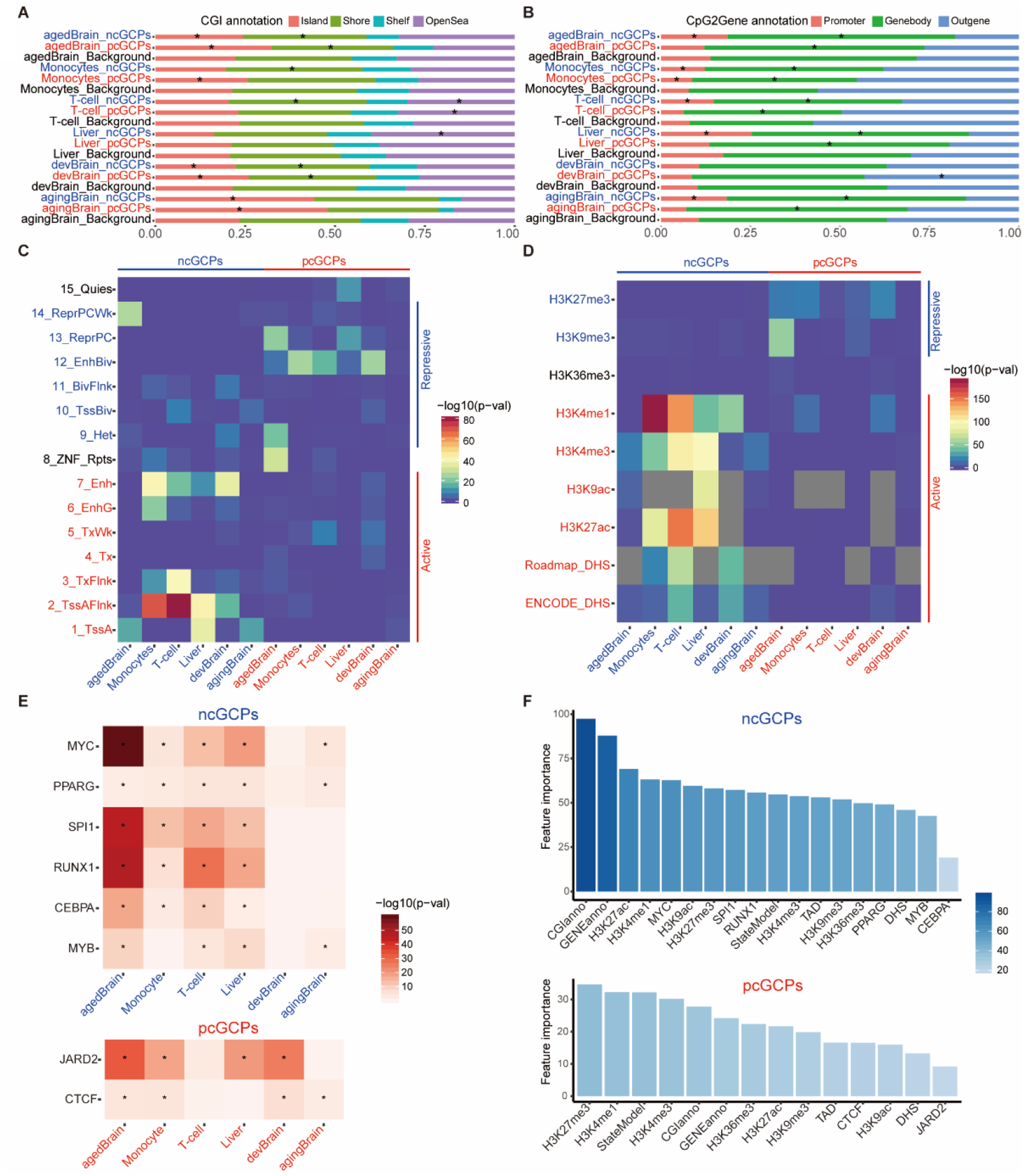
ncCpGs and pcCpGs are associated with multiple specific epigenomic features. (A) The bar plot for CpGs by location relative to CpG islands (CGI). The stars (*) represent significant enrichment in the hypergeometric test (p-value<0.05). agedBrain, brain aged group; devBrain, brain developmental group; agingBrain, brain aging group. (B) The bar plot for CpGs by location relative to gene body of the matched genes. The stars (*) represent significant enrichment in the hypergeometric test (p-value<0.05). (C) Overrepresentation of 15 epigenetic states in cGCPs. The active states are labeled as red color and the repressive states are labeled as blue color. (D) Overrepresentation of histone modification and DHS regions in cGCPs. The active markers are labeled as red and the repressive markers are labeled as blue. The markers which are not available are labeled as grey. (E) Overrepresentation of TF binding regions in ncGCPs and pcGCPs, respectively. The stars (*) represent significant enrichment in Fisher’s exact test (qvalue<0.1). (F) The feature importance for ncGCPs and pcGCPs, respectively.

Further, we evaluated the effects of the location of the cCpGs relative to their associated cGenes. pcCpGs and ncCpGs were found in different gene locations (**Fig. 6B**). Regardless of tissue specificity, the pcCpGs were more frequently located in the gene body while ncCpGs tended to be located on the promoter or gene body (p-value < 0.05). We also found that pcCpGs in the developmental brain were uniquely enriched in the outer regions of gene promoters and bodies (p-value < 2.2E-16), which was not observed in other tissues or brain of other age groups, suggesting a distinct regulatory mechanism in development.

In addition, we annotated the CpGs using the Roadmap chromatin state model and histone markers from matched tissues. The pcCpGs and ncCpGs possess different occurrence frequencies (p-value < 0.05) in multiple chromatin state regions and histone markers (**Fig. 6C, 6D**). The ncCpGs tended to be located in transcriptionally active regions and associated with actively epigenetic markers, such as histone acetylation and H3K4 methylation. In contrast, the pcCpGs were largely located in non-transcribed regions, especially in regions of repressed polyComb and bivalent enhancer, and associated with repressing markers like H3K27me3. Surprisingly, H3K4me1 as an active marker also showed enrichment on the pcGCPs.

The correlation direction can differ from one tissue to another because of the changed epigenetic environment. For example, cg03202060, cg24593918, and cg23464743 were all correlated with *HLA-DQB1* with negative correlations in the liver but positive correlations in T-cells, accompanied by different chromatin states and histone modification of these genomic regions in liver and T-cells (**Fig. 7**). Histone modifications and chromatin state differences could also contribute to the DNAm-expression correlation directionality of multiple CpGs correlated with the same gene in one tissue (**Supplemental Fig. S7-S10**). For example, in monocytes, *RPS16*-cg18487508 is a negative correlation while *RPS16*-cg20641794 is a positive correlation. Correspondingly, the cg18487508 and cg20641794 possess distinct chromatin states and histone modification in monocytes (**Supplemental Fig. S10**).

**Figure 7.**
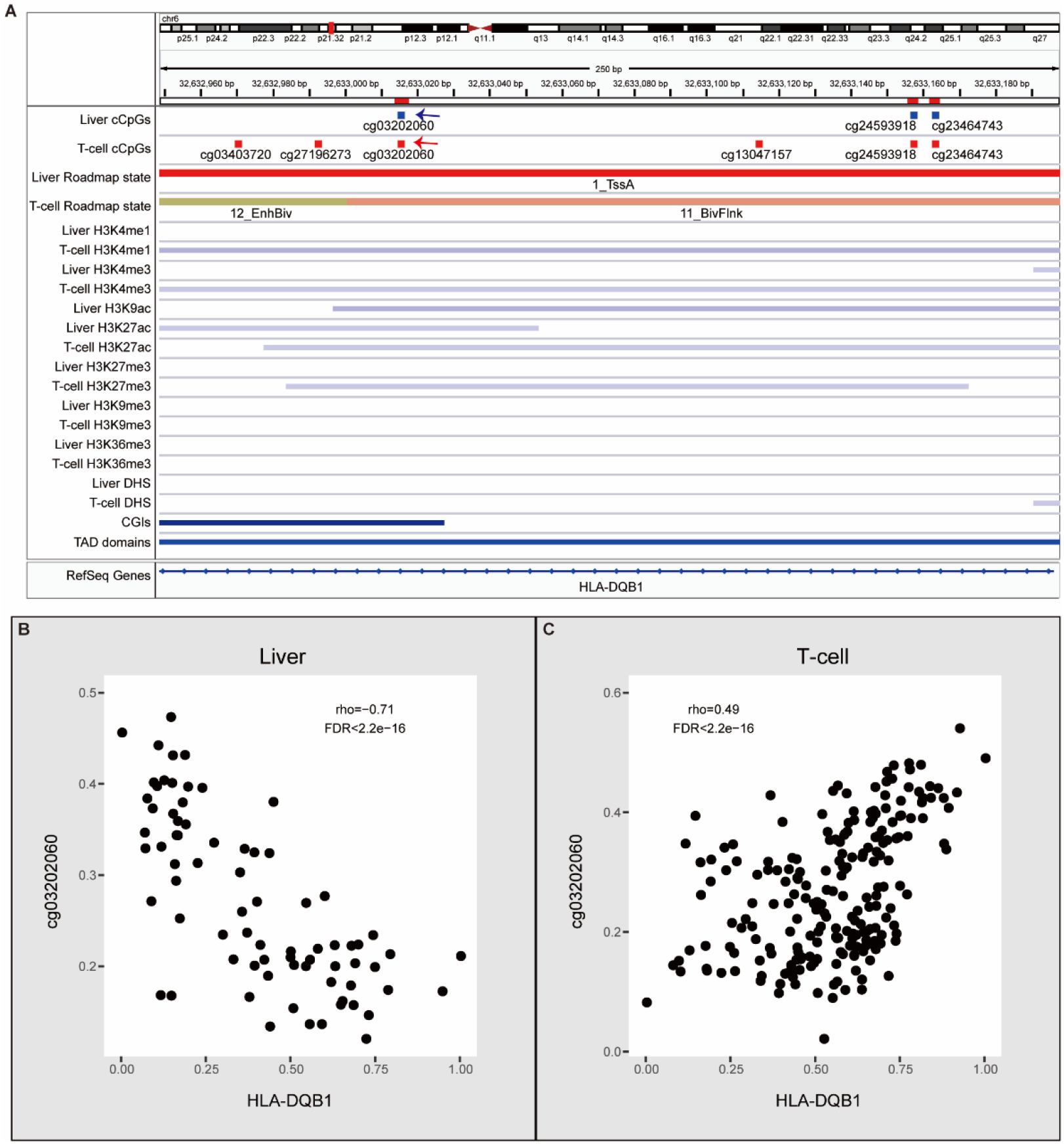
Examples of cGCPs with opposite correlations in liver or T-cells. (A) An IGV plot shows cCpGs of *HLA-DQB1* and epigenomic features. cCpGs in blue and red mean negative and positive correlations, respectively. The blue and red arrows point the examples of negative and positive cCpGs. (B) Scatter plots of *HLA-DQB1* expression level (X-axis) and cg03202060 methylation level (Y-axis) in the liver. (C) Scatter plots of HLA-DQB1 expression level (X-axis) and cg03202060 methylation level (Y-axis) in the T-cell.

To investigate the relationship between TFs, DNAm, and gene expression, we annotated the cCpGs using the human cistrome (the genome-wide map of regions bound by TFs). The ncCpGs were enriched in 6 TF binding regions in at least four of the six datasets, which included binding sites for MYC, PPARG, SPI1, RUNX1, CEBPA, and MYB (**Fig. 6E**). The pcCpGs were enriched in the JARD2 and CTCF binding regions in at least four of the six datasets (**Fig. 6E**). Some TFs-cCpG enrichments were observed only in specific tissue types, such as PO5F1-pcCpG in the developmental brain and FOXA1-ncCpG in liver only (**Supplemental Fig. S11**).

Topologically associated domains (TADs), a high level of 3D chromosome structure, are considered to mediate long-distance transcriptional regulation within their boundaries(Acemel et al. 2017). To confirm whether cGCPs are generally located within the same TAD, we downloaded the TAD regions of H1-hESC from ENCODE and found that most cCpGs and their corresponding cGenes (> 85%) were landed within the same TADs (**Supplemental Fig. S12**).

Finally, we determined the importance of epigenomic features in cGCPs using the random-forest method. We used the CpG location relative to CGI, gene body, chromatin state, histone modifications, TF binding sites, and the TADs mentioned above as input. The average decrease in accuracy measured by error rate was estimated after permuting each variable in ncGCPs and pcGCPs, respectively (**Fig. 6F**). As demonstrated in **Fig. 6F**, CpG location relative to CGI and gene body were found to be the two most important effectors for ncGCPs. Additionally, active histone modifications such as H3K27ac also have an important effect on the ncGCPs. The established gene repression marker, H3K27me3, was identified to be the most important effector for pcGCPs.

## Discussion

In this study, we evaluated DNAm and gene expression across different tissue types and brains of different age ranges, and found tissue- and age-specific DNAm-expression correlations. Thousands of GCPs were found to be correlated in various tissue types, but only a few pairs, primarily responsible for glutathione metabolic functions, were consistently correlated across different tissue types. DNAm-related regulation highlighted the functional importance of glutathione which is known to be critical in antioxidant defense, nutrient metabolism, and basic cellular functions(Wu et al. 2004; Forman et al. 2009). Moreover, glutathione level is directly related to methyl-donor and DNAm level(Lertratanangkoon et al. 1997). Thousands of cGCPs were identified in the developmental brain cohort but only four percent of these developmental cGCPs were also present in the aged brain. Most cGCPs in the developmental brain were found to be specific to neurogenesis. We further discovered new epigenetic signature for pcGCPs suggesting novel regulation mechanisms. Our findings suggest that DNA methylation patterns are essential for the maintenance of tissue- and development-specific gene expression.

This study confirmed the presence of both positive and negative relationships between DNAm and gene expression, and systematically assessed their associated epigenomic features. Our results support the well-known repressive model of DNAm in gene regulation by ncGCPs, including that DNAm at promoter and CGI regions represses gene expression(Jaenisch and Bird 2003), the involvement of active chromatin markers in negative cGCPs(Wagner et al. 2014), and the association with a large number of methylation-sensitive TFs(Maurano et al. 2015; Zhu et al. 2016). For the activator function of DNAm in regulating gene expression represented by pcGCPs, we discovered several associated epigenomic features. The DNAm activator model may be that increasing DNAm level could recruit the methylation-preferred activating TFs to promote gene expression. It is reported that some TFs, such as PO5F1 (*POU5F1*), preferred to bind the methylated DNA sequences(Yin et al. 2017). Another alternative model of DNAm activator may be modulated by the reduced binding of repressive TFs, like CTCF, which contributes to the establishment of the chromatin loops and genome topology(Ong and Corces 2014). DNAm could block the CTCF binding and open chromatin loop, which increased expression of the gene outside the loop(Liu et al. 2016). H3K27me3 regulated by PRC2 combined with JARD2 (*JARID2*) might be another example of pcGCPs (**Supplemental Fig. S14**). Our results showed the strong association of pcGCPs with the H3K27me3 marker regions and the JARD2. JARD2 recruits PRC2 to chromatin and may also regulate DNA methylation(Peng et al. 2009; Li et al. 2010; Pasini et al. 2010; Dixon et al. 2021). PRC2 is the only identified methyltransferase that catalyzes H3K27me3 and this process may be negatively regulated by DNAm(Bartke et al. 2010; Jermann et al. 2014; Wachter et al. 2014; Laugesen et al. 2019). In the H3K27me3 regions, increasing DNAm may inhibit the recruitment of PRC2 by JARD2 and prevent the deposition of H3K27me3, resulting in an open chromatin and gene activation. Our results suggested novel insight of DNAm-mediated expression regulation for the positively-associated GCPs.

Our study highlights the functional relevance of cGCPs by their enriched biological functions and human diseases. The cGCPs identified in our study were found to be associated with tissue- and age-specific disease phenotypes, suggesting an important contribution of methylation-correlated gene expression to complex traits. Based on our results, schizophrenia- and bipolar disorder-related cGCPs were preferentially enriched in developmental brain tissue, suggesting that those phenotypes are established during certain developmental stages. For example, *MDK* was only correlated with DNAm in the developmental brain but not in the aging and aged brains. *MDK* has been reported as a schizophrenia-related gene in several studies. It was identified as a schizophrenia-related gene by a *Pascal* gene-based test(Wu et al. 2017); a differentially expressed gene in brains of patients with schizophrenia compared to healthy controls(Gandal et al. 2018a); and a schizophrenia-associated gene in a transcriptome-wide association study (TWAS) in both adult brain(Gandal et al. 2018b) and fetal brain(Walker et al. 2019). The cGCPs associated with immune functions were also found to be linked to multiple psychiatric disorders, suggesting other important candidates for further investigation(Birnbaum et al. 2017; Hartwig et al. 2017). cCpGs related to lupus erythematosus were enriched in T-cells suggesting candidates for autoimmune disorders(Sharabi and Tsokos 2020). These results jointly imply that the concerted DNAm-expression are indicative to the biological functions of gene and CpG sites involved in the cGCPs.

We compiled a catalog of the correlated GCPs as references for other genetic and epigenetic studies (**Supplemental Table S2**). Genetic and transcriptome studies have yielded a large number of disease-associated candidate genes. DNAm abnormalities have also been identified in multiple diseases (Robertson 2005; Rakyan et al. 2011; Chen et al. 2014; Paul and Beck 2014; Li et al. 2019; Semick et al. 2019). With the cGCP catalog, we were able to connect disease-related genes with their putative DNAm regulators. It will reduce the search space for disease-relevant genes, a decrease in false positives, and increase statistical power.

Limitations in this study exist. First, the cGCPs identification is based on the correlation analysis which may be affected by factors, such as the sample size and variation of data(Goodwin and Leech 2006). Small sample size reduced the power in correlation tests and thus led to fewer significant cGCPs identified in these tissues such as the liver dataset. The regulatory relationships could be missed if both levels of DNAm and expression are constant in tissues. Correlation analysis also cannot decipher the causality. Though we have investigated the correlated DNAm-expression in large samples, more research is still needed to detect the correlated GCPs and interpret their causality in much more tissues and samples. Secondly, though we used the uniform preprocess pipeline, a batch factor may still confound the data from different laboratories and different platforms. Finally, since high-resolution Hi-C data was not available for the studied tissues, we used the H1-hESC TAD regions, which may not map the cGCPs interaction in all tissue precisely.

In summary, our study provides strong evidence that concerted DNAm-gene expression relationship is tissue- and development-specific. We also found that the positive and negative correlation direction can be switched in different tissues or age periods. We discovered novel epigenetic signature of pcGCPs, which suggested new DNAm-mediated regulatory mechanisms through CTCF and JARD2. Our findings highlight the functional importance of specific cGCPs for their cellular functions, and potential contribution to diverse disease susceptibility. Early developmental brain employed a highly unique set of cGCPs. *MDK* is one of the candidates of schizophrenia revealed by cGCPs. cGCPs identified in our study may facilitate further classification of regulatory elements and prioritization of tissue- and development-specific functional genes.

## Methods

### Data collection

To decipher the relationships between DNAm and gene expression in different tissues, we focused on the human tissues which had both methylation and matched expression data available in various databases and publications. Aged brain dorsolateral prefrontal cortex (DLPFC) data were obtained from the Rush University’s Religious Orders Study and Memory and Aging Project (ROSMAP) study (https://www.synapse.org/#!Synapse:syn3219045)(De Jager et al. 2014). Monocyte and T-cell data were both obtained from the MESA Epigenomics and Transcriptomics Study(Reynolds et al. 2014). Liver data were obtained from Horvath et al.(Horvath et al. 2014). To investigate the relationships between DNAm and gene expression across different ages, we used developmental and aging brain DLPFC data from Jaffe et al., with samples across the developmental stages ranging from prenatal to later postnatal life(Jaffe et al. 2016). The developmental brain was defined as stages from early fetal to 25 years old when brain development is complete(Pujol et al. 1993) and the aging brain was from 25 years old to aged stages. The data source, sample size, and platform for each dataset are summarized in **Supplemental Table S1**.

### Data preprocessing

DNA methylation measurements were performed using Illumina HumanMethylation450 BeadChips to interrogate more than 485,000 methylation sites in all datasets. Raw idat files were available for aged, developmental, and aging brain; raw intensity files were available for monocyte, T-cell, and liver. For methylation data analysis, we used the ChAMP package (Version1.8.2) implemented in R for filtering and normalization. We first defined the probe with a detection p-value more than 0.01 as an “absent” signal and the probe with bead number less than 3 as a “failed” probe. The samples with “absent” probes (more than 1% of total probes), the low-quality probes with “absent” signal (more than 10%), and “failed” probes (more than 5% across samples) were removed. The probes containing common SNPs as identified in Zhou et al.(Zhou et al. 2017) or mapped to multiple genome locations as identified in Nordlund et al.(Nordlund et al. 2013) were also discarded. Next, the beta value was used as the methylation measurement, and the beta-mixture quantile normalization (BMIQ) method(Teschendorff et al. 2013) was used to adjust the bias between type I and type II probes.

The expression data were collected from different platforms containing microarrays and RNA-seq. For the expression data from different platforms, we used different values as the expression measurement. Fragments per kilobase million (FPKM) was used to normalize RNA-seq data of the aged brain for correcting sequencing depth and the length of the gene. Genes were selected based on expression thresholds of >0.1 FPKM in at least 10% of samples for further analysis. For microarray expression data, filtering, normalization, and summarization were performed based on the platforms. For monocyte and T-cell data, we removed probes with a detection p-value above 0.06 in more than 20% of samples, and samples with more than 20% of absent probes were filtered out. The quantile normalized values were used for monocyte and T-cell data. For liver data, we used the R package *oligo* (Version1.36.1) to get the log2 RMA normalization as the gene-level expression values from CEL files. For developmental and aging brain data, the loess normalized log2 (sample/reference) ratios were used to quantify gene expression. Afterward, for the convenience of comparison, only protein-coding genes annotated in Gencode V19(Harrow et al. 2012) were kept.

For both expression and methylation datasets, several methods were used for quality control. Principal component analysis (PCA)(Martin and Maes 1979) and hierarchical cluster analysis were used to filter outliers of samples. ComBat(Johnson et al. 2007) was used to control for position and batch effects. Moreover, surrogate variable analysis (SVA)(Leek and Storey 2007) was applied to identify surrogate variables, and the confounding effects of known and unknown covariates were evaluated using principal variance component analysis (PVCA)(Li et al. 2009) and then regressed out on the data with the exception of age for the developmental and aging brain data. To remove the less variable methylation probes, we also filtered out 50% of methylation probes based on median absolute deviation (MAD).

### Statistical analysis of Gene and CpG Pairs (GCPs)

For each region, we defined pairs of methylation CpG site and gene expression as CpG sites located within the 10kb flanking region of the corresponding gene (**Fig. 1B**). Spearman’s rank correlation test was used to assess the correlation of methylation-expression pairs. Furthermore, we used the Benjamini-Hochberg (BH) method to correct for multiple hypothesis testing. We defined the significant threshold as a BH adjusted p-value <0.05.

### Cross- and downsampling-validation of cGCPs

To validate the power of the GCPs analysis, we randomly split each dataset in two-thirds percent as “discovery data” and one-third percent as “replication data”. We performed the GCPs correlation test in discovery data and replication data independently as described above. Then we calculated the Pearson correlation of the cGCPs’ rho value between discovery data and replication data. In total, we repeated the analysis one hundred times and found the Pearson correlation between discovery data and replication data to be very high (mean: 0.90~0.95, **Supplemental Fig. S1A, Supplemental Table S3**).

Because the sample sizes between each dataset differed (**Supplemental Table S1**), we randomly selected 75 individuals (the size of the liver dataset) from the aged brain, monocyte, and T-cell groups, and performed cGCPs correlation tests with these downsampled datasets to match the liver sample size. The Pearson correlation of the cGCPs’ rho value was highly correlated between the full and downsampled populations (mean: 0.86~0.96, **Supplemental Fig. S1B, Supplemental Table S3**).

### Functional annotation of cGCPs

For the genes of cGCPs, Gene Ontology (GO) enrichment was completed by the WEB-based GEne SeT AnaLysis Toolkit (WebGestalt)(Wang et al. 2017). Biological processes, cellular components, and molecular functions were tested using the overrepresentation enrichment analysis (ORA) method and genome protein-coding as a reference gene set. The BH method was used for multiple testing adjustment.

To explore the role of cGCPs in complex diseases and traits, we annotated the genes and CpGs of cGCPs using GWAS and EWAS results, respectively. For the genes of cGCPs, we performed the enrichment analysis of GWAS catalog reported gene sets using FUMA GWAS v1.3.6a(Watanabe et al. 2017). The minimum overlapping genes with gene-sets was set to 2. Protein-coding genes were used as background genes and the BH method was used for multiple testing corrections. For the CpGs of GCPs, we downloaded the EWAS results (updated 03-07-2019) from the EWAS catalog (http://www.ewascatalog.org/). We used the results with p < 2.4×10-7 for the 450k array(Saffari et al. 2018). We tested the CpG sets with a minimum number of overlapped CpGs to 2. Hypergeometric tests were performed to test if the CpGs were overrepresented in any of the EWAS CpG sets. Significant enrichment was defined as BH adjusted p-value < 0.05.

### Epigenomic features of cGCPs

The CpGs and genes involved in cGCPs were annotated using multiple epigenomic features. CpGs of cGCPs were defined accordingly: 1) by location relative to CpG Islands, CpGs were classified as CpG island (CGI), CGI shores (0-2,000bp up- or down-stream of CGI), CGI shelves (2,000-4,000bp up- or down-stream of CGI), and open sea region (>4,000bp up- or down-stream away from CGI); 2) by location relative to the matched gene, CpGs were classified as gene promoter (2kb upstream to transcription start sites (TSS) defined in Gencode v.19), gene body region (TSS to transcription end site (TES)), outgene region (out of promoter and gene body). Fold enrichment was tested against the total tested GCPs in each tissue. The significance of enrichment was determined by the hypergeometric test.

To assess CpGs for epigenetic state enrichment, we downloaded the histone marks and DNaseI hypersensitivity sites (DHS) data of the matched tissues from the Roadmap Epigenomics Project (http://www.roadmapepigenomics.org/) and the ENCODE Project (https://www.encodeproject.org/) (**Supplemental Table S20**). In addition, the 15-epigenetic state model of the matched tissue from the Roadmap Epigenomics Project was used to generate the epigenetic state table (**Supplemental Table S20**). Using the CpGs of total GCPs analyzed in each tissue as a background, we performed the epigenetic state enrichment of the CpGs from cGCPs using the R LOLA package(Sheffield and Bock 2015).

We downloaded the Hi-C defined topological domains (TAD) of H1-hESC for GCPs annotation. The data were downloaded from Dixon et al.(Dixon et al. 2012) and the coordinates were converted to hg19 using UCSC’s liftOver tool by Ho et al.(Ho et al. 2014). The cGCPs within the same TAD were annotated using bedtools (v2.17.0)(Quinlan and Hall 2010).

To test the effect of transcription factors (TFs) on cGCPs, we downloaded the human cistromes which provide the genome-wide maps of regions bound by TFs from Vorontsov et al.(Vorontsov et al. 2018). To achieve the highest reliability, we only used the cistrome regions detected in at least two experimental data sets and by at least two peak calling tools. A total number of 138 TFs were tested using the R LOLA package(Sheffield and Bock 2015). Q-value in R *qvalue* package was used to controlling for the false discovery rate. Significant TF enrichment was defined as q-value < 0.1.

To estimate the importance of functional features on cGCPs, the random forest algorithm, implemented by the R package *“randomForest”*, was applied to pcGCPs and ncGCPs, respectively. Feature importance was tested on the basis of the average decrease of accuracy by permuting each variable.

## ACKNOWLEDGMENTS

This work was supported by the National Natural Science Foundation of China (Grants Nos. 82022024, 31970572, 31871276), the National Key R&D Project of China (Grants No. 2016YFC1306000 and 2017YFC0908701), Innovation-driven Project of Central South University (Grant Nos. 2020CX003), and NIH grants U01 MH122591, 1U01MH116489, 1R01MH110920. We sincerely thank Liz Kuney, Richard F. Kopp, and other colleagues for scientific and language advice that helped improve our manuscript.

## Author Contributions

KW, RD, YX, and CJ collected and preprocessed the datasets. KW performed the GCPs analysis. KW and JH performed the feature and GWAS enrichment analysis. KW, CC, TM, CZ, and CL participated in the revision of the manuscript. CC and CL conceived the study, participated in its design, and supervised the entire project. All authors read and approved the final manuscript.

